# Efficient planning and implementation of optimal foraging strategies under energetic constraints

**DOI:** 10.1101/2025.07.04.663150

**Authors:** Yipei Guo, Ann M Hermundstad

## Abstract

To successfully forage for food, animals must balance the energetic cost of searching for food sources with the energetic benefit of exploiting those sources. While the Marginal Value Theorem provides one normative account of this balance by specifying that a forager should leave a food patch when its energetic yield falls below the average yield of other patches in the environment, it assumes the presence of other readily reachable patches. In natural settings, however, a forager does not know whether it will encounter additional food patches, and it must balance potential energetic costs and benefits accordingly. Upon first encountering a patch of food, it faces a decision of whether and when to leave the patch in search of better options, and when to return if no better options are found. Here, we explore how a forager should structure its search for new food patches when the existence of those patches is unknown, and when searching for those patches requires energy that can only be harvested from a single known food patch. We identify conditions under which it is more favorable to explore the environment in several successive trips rather than in a single long exploration, and we show how the optimal sequence of trips depends on the forager’s beliefs about the distribution and nutritional content of food patches in the environment. This optimal strategy is well approximated by a local decision that can be implemented by a simple neural circuit architecture. Together, this work highlights how energetic constraints and prior beliefs shape optimal foraging strategies, and how such strategies can be approximated by simple neural networks that implement local decision rules.

## Introduction

Searching for new food sources is energetically costly. Successful foragers must balance this energetic cost against the gains of exploiting known food sources. The Marginal Value Theorem (MVT) provides a normative framework for describing this trade-off [1, 2]. In its original formulation, food is assumed to be distributed across discrete patches, and travel between patches incurs a fixed time cost. A forager gains energy at a diminishing rate while exploiting a patch due to resource depletion. The optimal policy, derived by maximizing the long-term average rate of energy intake, prescribes that a forager should leave a patch when its energetic yield falls below the average yield of other patches in the environment [1]. Crucially, this approach assumes that the forager will reliably encounter another patch after each departure.

This basic framework has been extended in numerous ways to account for additional ecological and behavioral factors. Some models consider uncertainty about patch quality [3, 4], or incorporate movement decisions such as how fast to travel between patches [5]. Other studies relax the assumption of fixed travel times by modelling inter-patch search as a stochastic search process [6, 7], and such mechanistic models have been used to examine how the type of random walk affects patch-finding success [8, 9]. Yet, across all these model extensions, a key assumption typically remains: that other food patches exist and will eventually be found.

In natural environments, however, the existence of additional food sources is often unknown, and a forager must weigh the energetic costs of unsuccessful search against the potential benefits of discovering new patches. Many insects, including flies, have been observed to engage in local exploratory search around a known food source [10–12], and the average duration of their exploration increases over time [13]. This strategy—of first sampling the nearby environment before expanding outward if no alternatives are found—may be an efficient way to conserve energy while maintaining access to a reliable food source. Alternative strategies—such as beginning with a more extensive exploration of the environment in the hope of quickly identifying better food sources—carry a higher risk of energetic depletion, but may offer long-term benefits by enabling more efficient foraging later on. These different possibilities raise a fundamental question about optimal foraging: upon first encountering a patch of food, how should a forager structure its search for new patches when (i) the existence of those patches is not known with certainty, and (ii) searching for those patches requires energy that can only be harvested from a single known food patch?

Here, we explore optimal strategies for structuring search under such energetic constraints. Because the forager faces the possibility that there will not be any food patches nearby, we assume that it initiates a search only when, and for as long as, it has enough energy to reliably return to a known food patch. In these settings, we show that it can be advantageous to partition the search into multiple successive trips, depending on the forager’s belief about the distribution and content of patches in the environment. Although the optimal search strategy often requires planning multiple trips into the future, we find that a local decision rule can closely approximate optimal performance and is implementable in a biologically plausible circuit architecture. Together, this work provides insights into how foragers should best balance energetic costs and demands when searching for food in uncertain environments, and how this energetic balance can be achieved at the level of neural circuits and behavior.

## Results

### General setup

We consider a hungry forager that has just discovered a patch of food in a new environment (Fig 1a). After harvesting some food from the patch, the forager must decide whether to continue harvesting from the same patch, or whether to leave the patch and seek out better options. If there are no additional patches in its vicinity, the forager might be best served by harvesting as much as possible from the patch in order to gain enough energy to make a long trip to another environment. However, if there are nearby patches with higher yields or complementary resources, the forager would benefit from more quickly seeking out and exploiting these nearby patches.

**Figure 1.**
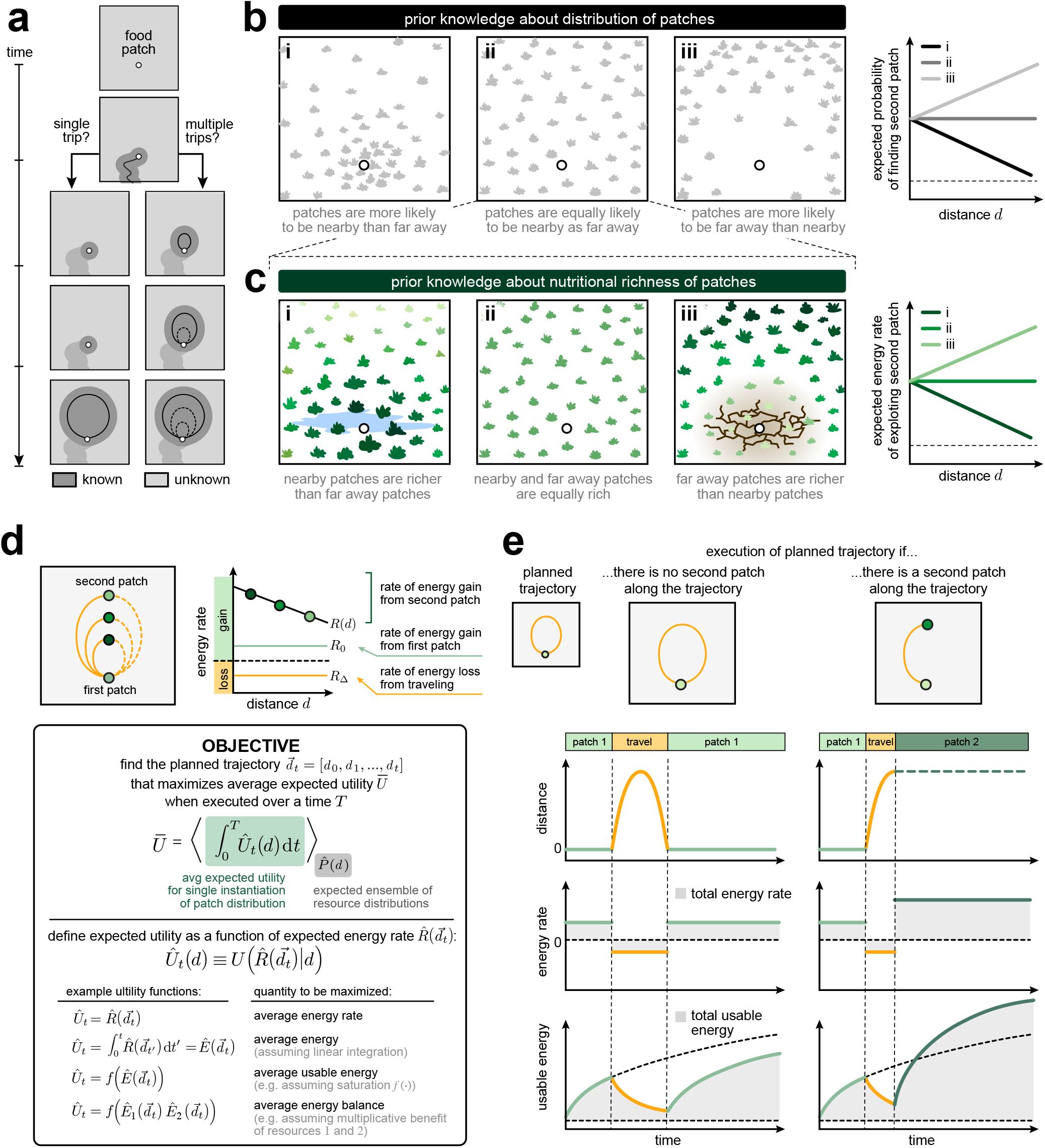
An energetically-constrained forager faces a decision of how often and how far to travel. **a)** A forager on a food patch must decide whether to explore and in how many trips. **b-c)** The forager maintains knowledge about the spatial distribution (b) and nutritional content (c) of food patches. **d)** We consider a 1D scenario in which the forager gains energy at a rate *R*_0_ while harvesting from a known food patch, loses energy at a rate *R*_Δ_ while traveling, and can gain additional energy at a rate *R*(*d*) if a second food patch is found at a distance *d* from the original food spot. Box: To weigh the potential costs and benefits of traveling away from the first food patch, we measure the average expected utility 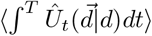 of executing a planned trajectory 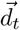, averaged over all possible locations of the closest food patch *d. U* can take on many forms, depending on the specific demands of the forager. **e)** The forager seeks the trajectory that maximizes the average expected utility over time and across possible distributions of patches. In planning this trajectory, the forager must account for the possibilities that it does (right column) or does not (left column) encounter a second patch along its trajectory.

Without knowing which of these situations it faces, the forager must use its prior belief about the distribution and nutritional content of food patches in the environment to decide whether it is beneficial to search, and if beneficial, how far to travel. Because the forager faces the possibility that there will not be any food patches nearby, we assume that it structures its search in a series of outbound and inbound trips to ensure that it returns to the known food patch if it fails to find a second patch. The forager must then decide whether to search the environment in a single long trip, or whether to break the search into multiple trips (Fig 1a). The optimal number and sequence of such trips will depend on the forager’s prior beliefs about the environment, and it will be constrained by how much energy the forager can harvest from the original food spot.

To guide its decisions about whether and when to leave the first food patch, we assume that the forager maintains prior expectations about the distribution of food patches that it is likely to encounter in a new environment (Fig 1b) and the nutritional content of those food patches (Fig 1c). We assume that knowledge about these two environmental features is maintained in two separate functions that depend only on distance from the forager: (i) the expected probability 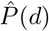 of finding a second food patch at a distance *d* away, and if found, (ii) the expected rate of energetic return per unit time, 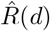, of the second patch. The shape of these two functions determines the relative probability and energetic return of finding food patches that are nearby versus far away. For example, if the expected probability of finding a second food patch decays with distance, the forager expects that the first food patch is indicative that other food patches are more likely to be clustered nearby rather than far away. Alternatively, if the expected energy rate of a second food patch decays with distance, the forager expects that nearby food patches will either have higher nutritional yield or will be more nutritionally diverse than distant patches. As we will show, the interaction between these two functions will shape the forager’s optimal search strategy. In what follows, we will assume that both 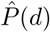 and 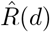 reflect the true statistics of the environment *P* (*d*) and *R*(*d*) (i.e., the forager maintains the correct generative model of food patches in the environment).

In order to decide whether to make an exploratory trip, the forager must weigh the expected gain in energy from finding a second patch along the outbound trip (minus the expected cost of outbound travel) against the expected cost of traveling the full outbound and inbound trip, should there be no second patch (Fig 1d). We assume that the forager harvests energy at a fixed rate *R*_0_ *>* 0 while on the first food patch and expends energy at a fixed rate *R*_Δ_ *<* 0 while traveling, and we further assume that the values of *R*_0_ and *R*_Δ_ are known to the forager. To weigh the potential costs and benefits of traveling away from the first patch, we assume that the forager plans a trajectory consisting of one or more successive outbound and inbound trips, subject to the constraint that the forager’s energy cannot drop below zero at any point along the trajectory. To compare different planned trajectories, we evaluate the expected utility of each trajectory over a total time *T*. Because the forager only maintains knowledge about the expected *distribution* of possible patches 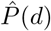, but not about the specific instantiation of that distribution, we compute the average expected utility over all possible instantiations of 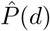. For each instantiation, we assume that the forager would carry out its planned trajectory unless or until it encounters a second patch, at which point it would harvest energy at an average rate *R*(*d*^*\**^) that is determined by the location *d*^*\**^ of the second patch (in this way, *R*(*d*) can represent the instantaneous energy rate of the second patch or the average rate of traveling between and exploiting two complementary patches). We define the expected utility 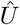 to be a function of the expected energy rate 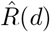. Different functional forms of 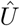 correspond to different quantities that the forager might aim to maximize, such as the average rate or amount of usable energy harvested from a single patch, or the average energetic balance between different types of patches (Fig 1d).

In what follows, we explore how the forager’s expectations about the distribution and quality of food patches impacts its planned foraging trajectory. For tractability, we consider a simplified one-dimensional scenario in which the forager travels at a constant velocity *v* = 1 when traveling away from and returning back to a known food patch at the origin, *d* = 0, within an environment of size *D* (Fig 1e). In this case, a planned trajectory consists of a sequence 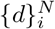 of *N* outbound trips to and return trips from locations *d*_1_, *d*_2_…*d*_*N*_ initiated at times *t*_1_, *t*_2_, …*t*_*N*_ . We take food patches to be distributed with fixed probability *P* (*d*) = *p*, such that the location of the nearest patch follows *P*_1_(*d*) = *pe*^*−pd*^. When the forager encounters a second patch, we assume that it can harvest energy at a rate *R*(*d*) = *µ* (1 *− γd*), with *µ > R*_0_ (such that there is a potential benefit to exploration) and *γ ≥* 0 (such that there is no benefit from further exploration once a second patch is found). We evaluate trajectories with respect to the utility function *U* = *R* (SI section 1), which corresponds to maximizing the average energy rate over a time interval *T* (or, equivalently, the final energy at time *T* ). Under this assumption, for a planned outbound trip to location *d*, the forager should leave the food patch as soon as it has enough energy to travel the outbound and return distance 2*d*, such that the time spent on the food patch before a trip to *d* is given by Ω(*d*) = 2*d*|*R*_Δ_|*/R*_0_.

### Moderately dense and nutritionally rich environments encourage multiple trips

We began by exploring whether, and under what conditions, it would be better for the forager to plan to explore the environment in two successive trips versus one long trip. We first determined the optimal single trip to and from a location *d*_*L*_ that would maximize the expected utility over a time interval *T*, given a fixed set of environmental parameters *p, γ*, and *µ*. We then determined the scenarios under which it would have been favorable to first plan a shorter trip to and from a location *d*_*S*_.

The change in expected utility from having an additional first trip, Δ*U*, depends on the forager’s expectation of encountering a second food patch. This expectation can be decomposed into three different scenarios that occur with different probabilities, depending on the density of patches *p*:

#### Scenario 1: No second patch

If there is no second patch within a distance *d*_*L*_, it would be detrimental to first take a short trip because the forager would lose additional energy (*R*_0_ *− R*_Δ_) while traveling the distance 2*d*_*S*_ (relative to using the same time to harvest from the first patch at a rate *R*_0_). This scenario occurs with probability *e*^*−pd*^*L*, and leads to an expected *loss* in utility (Fig 2b, left column):

**Figure 2.**
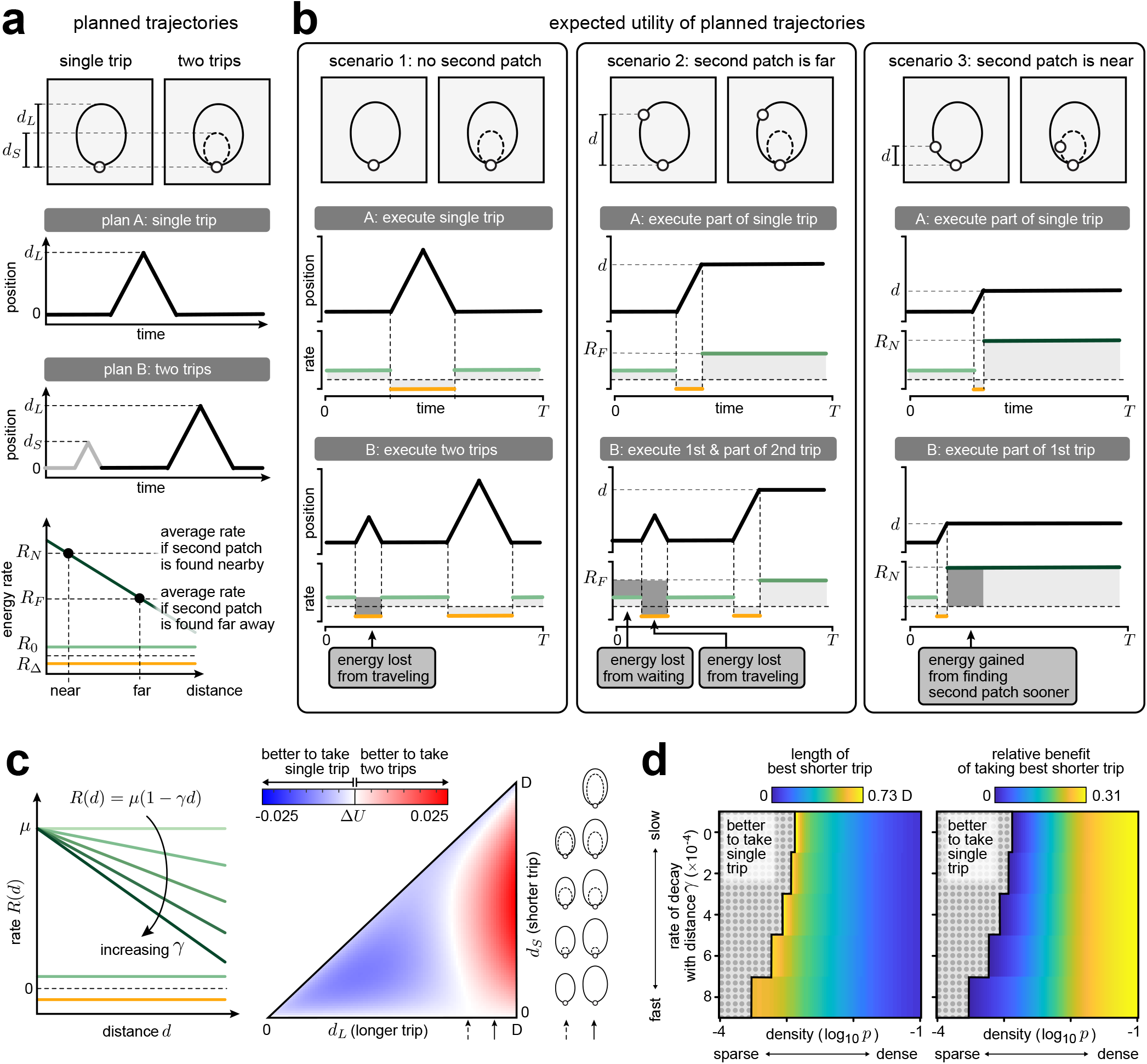
In certain environments, it can be advantageous to explore in multiple trips. **a)** Given a single trip of length *d*_*L*_, we evaluate the relative gain in utility of first taking a shorter trip of length *d*_*S*_. **b)** The relative gain of taking a short trip before a long trip can be decomposed into three different contributions: (i) the energy lost from the short trip if no second patch is found (left column), (ii) the energy lost from waiting and traveling if the second patch is beyond the reach of the shorter trip (middle column), and (iii) the energy gained from finding the second patch sooner if it is within reach of the second trip (right column). **c)** We use *γ* to parameterize the rate of energy gain *R*(*d*) from patches a distance *d* away. For a given value of *γ*, we can determine the relative gain in utility Δ*U* of taking a short trip of length *d*_*S*_ before a longer trip of length *d*_*L*_. We select the best combination of short and long trips for which Δ*U >* 0; if Δ*U <* 0, the agent should use a single trip to explore the environment. **d)** As *p* increases and as *R* decays more rapidly with distance (i.e., *γ* increases), it becomes more beneficial (right) to take an increasing short second trip (left). [Parameters: (c-d) *D* = 1000, *R*_0_ = 0.1, *R*_Δ_ = *−* 0.1, *µ* = 10, *T* = 8000, (c) *γ* = 0.001, *p* = 0.24771.]

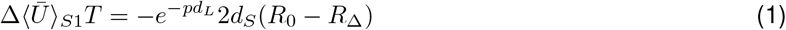

#### Scenario 2: Second patch is far

If there is a second patch at a distance *d > d*_*S*_, it would again be detrimental to first take a short trip. This is not only because the forager would lose additional energy (*R*(*d*) *− R*_Δ_) while traveling the distance 2*d*_*S*_ (relative to using the same time to harvest from the second patch at a rate *R*(*d*)), but also because in preparing to take the first trip, the forager would lose additional energy Ω(*d*_*S*_)(*R*(*d*) *− R*_0_) as a result of harvesting from the first patch (relative to using the same time to harvest from the second patch at a rate *R*(*d*)). When averaged over all possible locations *d*_*S*_ *< d < d*_*L*_ of the second patch, each occurring with probability *P*_1_(*d*), this leads to an expected *loss* in utility (Fig 2b, middle column):

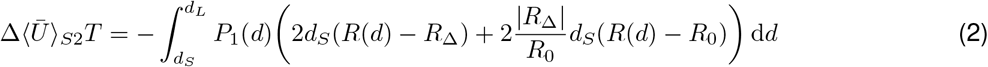

#### Scenario 3: Second patch is near

If there is a second patch within a distance *d*_*S*_, it would be beneficial to first take a short trip because the forager could find and exploit the second patch more quickly, thereby gaining additional energy (Ω(*d*_*L*_) *−*Ω(*d*_*S*_)) (*R*(*d*) *− R*_0_) (relative to using the same time to harvest from the first patch at a rate *R*_0_). When averaged over all possible locations *d < d*_*S*_ of the second patch, each occurring with probability *P*_1_(*d*), this leads to an expected *gain* in utility (Fig 2b, right column):

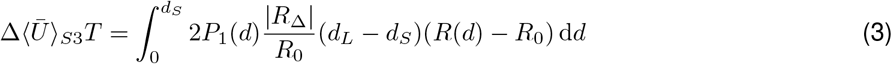

When planning a sequence of trips, the forager does not know which of these three scenarios it will face, and thus all three scenarios contribute to the expected benefit of exploring the environment in two trips rather than one Δ*U* = Δ ⟨ *Ū* ⟩*Ū*+ Δ ⟨*Ū* ⟩*Ū*+ Δ ⟨*Ū*⟩ *Ū*(Fig 2b). If the expected nutritional content is constant across space (i.e., *γ* = 0 and 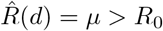), and in the limit that patches are sparsely distributed (*pD«* 1), the expected loss in utility from Scenario 2 always exceeds the expected gain in utility from Scenario 3, regardless of the lengths of the two trips (SI Section 2). As a result, it is always favorable for the forager to explore the environment in a single trip (Fig 2d; upper left of each heatmap). As the density of patches increases, there is a greater probability of finding food close to the original patch. This reduces the expected loss in utility from Scenario 1, and increases the expected gain in utility from Scenario 3, thereby encouraging the planning of two trips over one. Similarly, as the expected nutritional content decays more sharply with distance (i.e., as *γ* increases above 0), the expected gain in utility from Scenario 3 begins to increasingly outweigh the expected loss from Scenario 2, again encouraging the planning of two trips over one.

Together, these results show that it becomes increasing beneficial to break a single exploratory trip into a sequence of two trips in environments with a higher density of patches, and in environments where nearby patches are expected to be more nutrient-rich than distant patches (whether because they have higher yield, or whether because it requires less energy to travel between and exploit multiple patches simultaneously).

### Planning for multiple trips can further improve performance

The previous results identified environmental conditions under which it is favorable for the forager to structure its planned exploration in two successive trips rather than one. This suggests that increasing the planning horizon beyond two trips might confer an additional benefit.

To explore how expected utility depends on the planning horizon *M*, we numerically solved for the optimal sequence of *K* planned trips 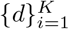, with 0 *< d*_1_ *< d*_2_… *< d*_*K*_ *≤ D*, for all values of *K* = 1, 2, 3, …, *M* (SI section 1). We denote *K*_opt_ *≤M* as the optimal value of *K* that maximizes expected utility (note that in the limit of large *M*, the optimal sequence of *K*_opt_ trips is the globally optimal solution, and if *K*_opt_ *< M*, increasing *M* does not change the optimal solution). Consistent with the results discussed in the previous section, we find that the optimal number of trips *K*_opt_, and thus the optimal planning horizon, increases as the density of resources increases (Fig 3a-i). In other words, resource-rich environments encourage the planning of multiple exploratory trips.

**Figure 3.**
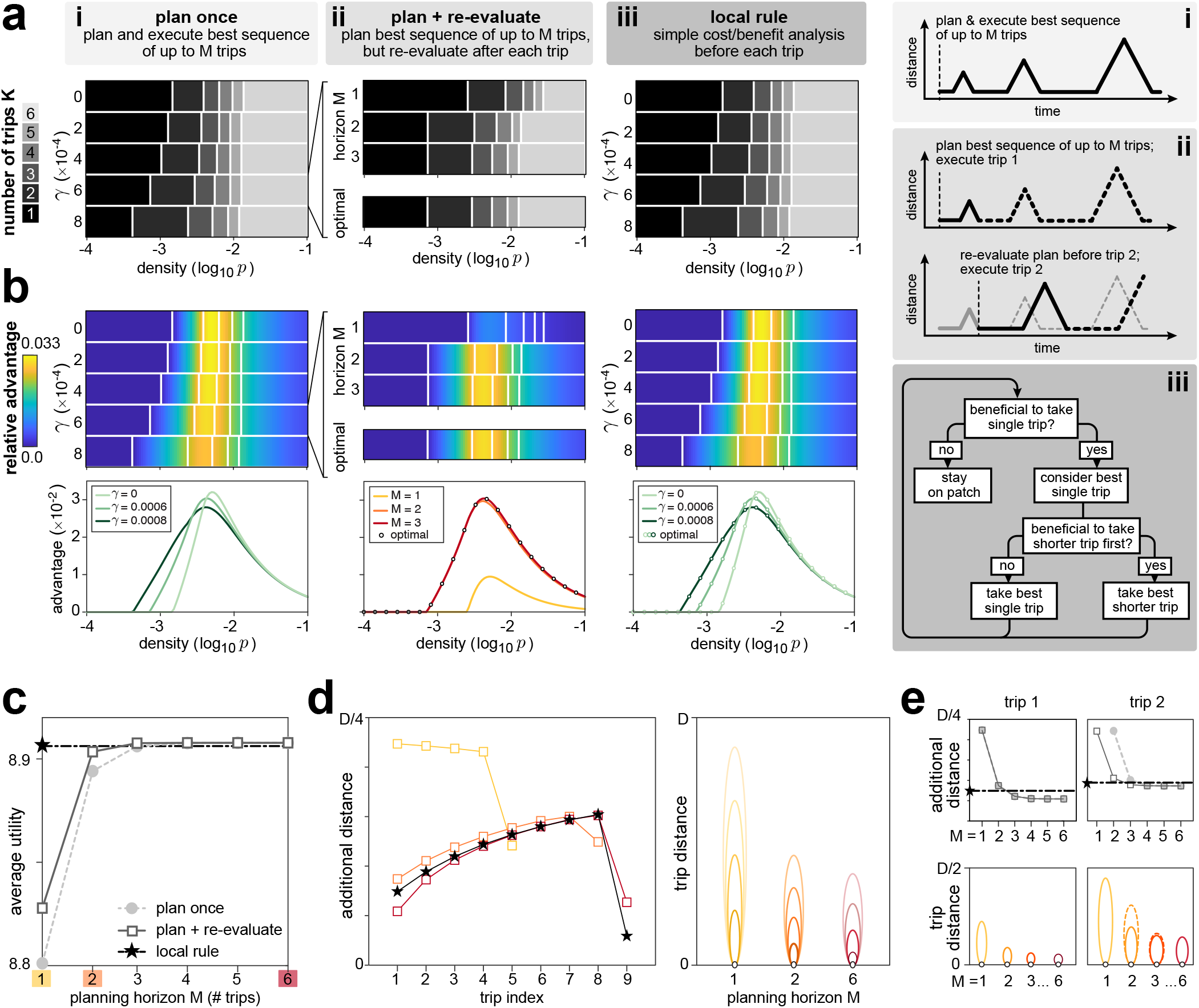
Planning for multiple trips can improve performance. **a)** Optimal number of trips for different environmental conditions, compared across three different strategies: (i) planning and executing a single optimal trajectory without re-evaluation, (ii) re-evaluating the optimal trajectory after each successive trip, and (iii) implementing a local decision rule on each trip. Strategies (i)-(ii) impose a maximum planning horizon *M*, and determine the best sequence of *K ≤ M* trips below that planning horizon. For strategy (i), the planning horizon must equal or exceed the optimal number of trips in order to achieve optimal performance. For strategy (ii), the optimal number of trips can be achieved with a shorter planning horizon. Strategy (iii) achieves optimal performance with a local rule (analogous to a planning horizon of *M* = 1). **b)** Same as (a), but evaluating the relative advantage (Eq. 4) of taking the optimal sequence of *K* trips compared to best single trip. **c)** Average utility as a function of planning horizon *M* . Longer planning horizons lead to higher utility; however, the local decision rule can achieve near-optimal utility with only local operations. **d)** Left: additional distance covered by successive trips for different strategies highlighted in **c**. Right: illustration of successive trips highlighted in left panel, split out by different planning horizons. In general, longer planning horizons lead to shorter initial trips; allowing for re-evaluation further changes subsequent trip lengths. **e)** Additional distance (upper) and trip distance (lower) as a function of planning horizon, shown for first and second trips. [Parameters: (a-e) *D* = 1000, *R*_0_ = 0.25, *R*_Δ_ = *−*0.25, *µ* = 10, *T* = 24000, (c-e) *p* = 0.24771, *γ* = 0.0006.]

We next measured the relative advantage *s*_opt_ of planning and executing the optimal sequence of *K* trips over that of executing the optimal single trip:

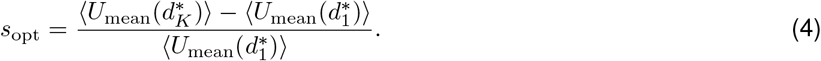

By computing this relative advantage over a range of environmental conditions, we find that planning for multiple trips confers the greatest advantage when resources are moderately dense (Fig 3b-i). When the density is sufficiently low, the forager is best served by planning a single long trip (as in Fig 2d). Alternatively, when the density is sufficiently high, the expected utility of a single trip is high, and thus the forager gains little by planning any additional trips.

For a given set of environmental conditions, the average utility increases as the planning horizon *M* increases (Fig 3c), and approaches the globally-optimal solution for large *M* . By comparing the planned distances of each trip 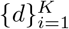 for different planning horizons *M*, we also find that *d*_*i*_(*M* ) decreases with *M* for *i ≤ K ≤ K*_opt_, implying that planning for the possibility of taking multiple trips decreases the planned length of each trip (Fig 3d).

### Re-evaluating after every trip can improve performance, but still requires planning for multiple trips

The previous results assume that the forager carries out a planned sequence of trips until it encounters a new food patch, without re-evaluating that plan based on newly acquired information about the environment (namely that explored regions do not contain food). We next asked whether re-evaluation could reduce the optimal planning horizon. To this end, we allowed the forager to re-evaluate a planned sequence of trips after each successive trip (note that this only occurs when the forager did not encounter a patch during the previous trip). As before, we assume that the forager determines the best sequence of trips up to a planning horizon of *M* trips, but we now consider the case where the forager takes only the first trip in that sequence. If, on that trip, the forager encounters a food patch at a position *d*^*\**^, we assume as before that the forager harvests energy at an average rate *R*(*d*^*\**^) for the remainder of the time *T* . If no new food patch is encountered, the forager returns to the original food patch, updates its belief about the availability of food sources, and plans a new sequence of up to *M* trips. The simulation stops when the time exceeds *T*, or when it is no longer favorable to explore (which can occur, for example, when the animal has already explored the whole space, or when the remaining unexplored space provides a lower expected energy rate than the original food patch). After each trip, the forager’s belief about the probability of encountering new food patches is updated to:

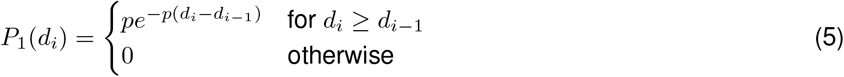

For relatively short planning horizons, this strategy quickly achieves optimal performance, even when the optimal number of trips exceeds the limit of the planning horizon (Fig 3a-ii, b-ii). Moreover, for a given planning horizon *M*, planning with re-evaluation enables the forager to achieve a higher average utility than planning without re-evaluation (Fig 3c); equivalently, with re-evaluation, the forager can achieve the same desired utility with a shorter planning horizon. Nevertheless, even with re-evaluation, planning for multiple trips can significantly increase expected utility (Fig 3c).

One reason that re-evaluation improves performance is that it relaxes the constraint that we otherwise impose on the maximum number of trips that can be carried out in any fixed period of time; in other words, because the forager can re-plan after each trip, the total number of trips is not limited by the planning horizon. This is illustrated in Fig 3c-d, where the ability to re-evaluate separates the behavior and performance of two foragers with a limited planning horizon of *M* = 1. Both foragers take the same first trip (Fig 3d, lower left), but the forager that re-evaluates takes a second trip (Fig 3d, lower right), which increases its average utility (Fig 3c). However, the process of re-evaluation still benefits from a longer planning horizon, since the possibility of taking multiple trips can be important for deciding the optimal length of each subsequent trip.

### A local decision strategy can achieve close-to-optimal performance

While iterative planning for multiple trips can improve overall performance, it is computationally costly. We thus asked whether a simpler strategy could achieve comparable performance. To this end, we constructed a two-step local rule for determining whether to leave the food patch, and how far to travel on each successive trip:

**Step 1:** Determine the optimal length of a single trip,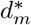. If the expected utility of this trip is greater than that of staying on the original food patch, proceed to Step 2; otherwise, continue harvesting from the original patch.

**Step 2:** Weigh the expected benefit and cost of first taking a shorter trip to 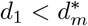, and determine the optimal value of 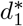 that maximizes the benefit minus the cost. If the benefit of taking 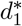 exceeds the cost, plan to travel to 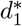; otherwise, plan to travel to 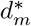.

Through simulation, we find that this simple local rule achieves near-optimal performance (Fig 3a-c).

### The local decision strategy can be implemented in simple neural network architecture

Having shown how optimal decisions can be achieved by a simple local decision rule, we now show how this decision rule could be implemented within a simple neural network architecture. This architecture consists of two parallel but interacting streams of operations that are carried out over several feedforward layers with skip connections (Fig 4a):

**Figure 4.**
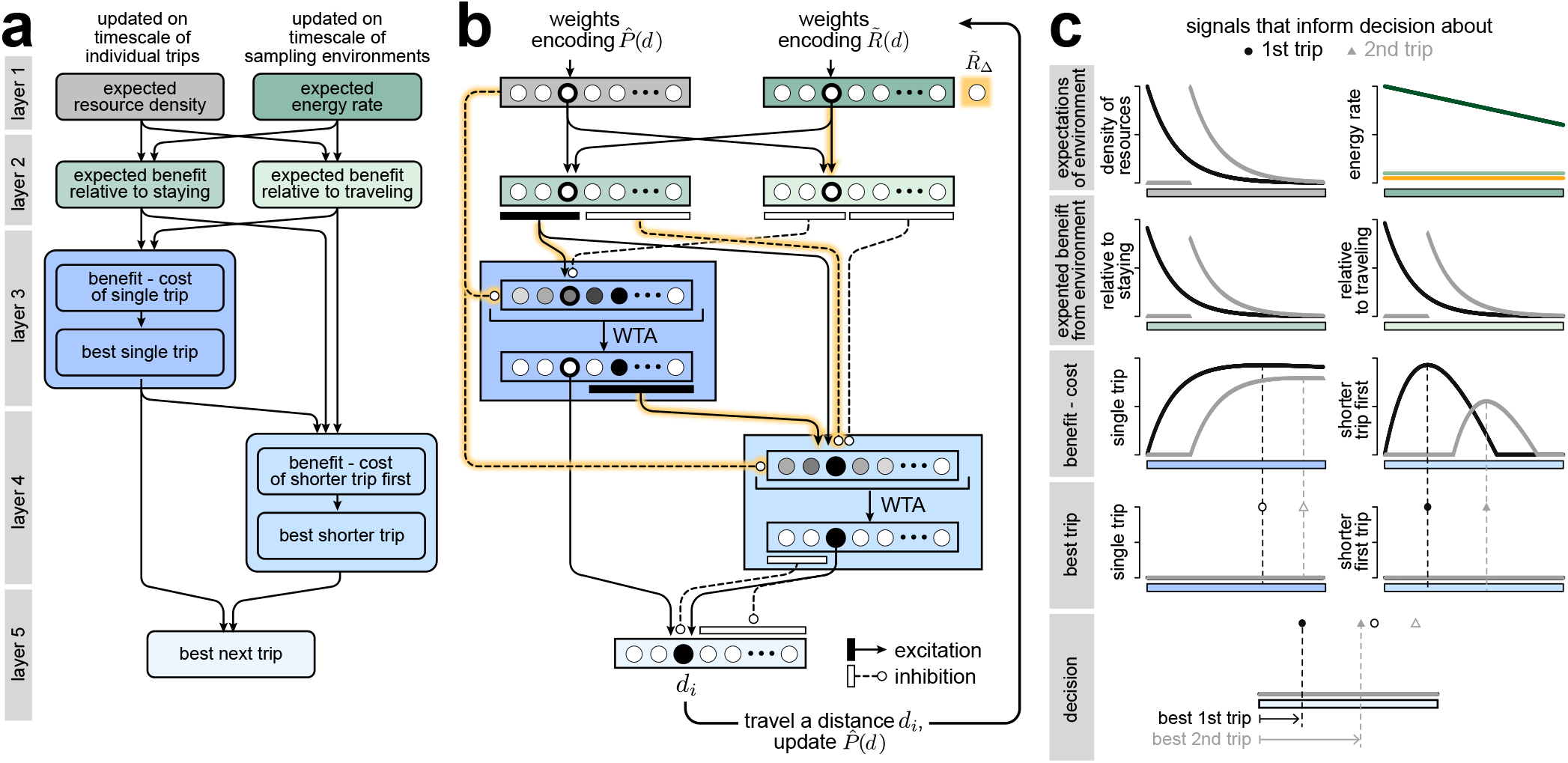
A simple neural network can implement the local decision rule. **a)** The local decision rule can be broked down into a set of successive computations that are performed in different modules of a neural network, as described in the text and as elaborated in panel **b. b)** A network architecture can implement the local rule using different modules of neurons (colored boxes), each ordered by distance. Weights between modules can be derived from the expressions for the local decision rule, and involve multiple stages of excitatory and inhibitory connections between successive layers. A neuromodulatory signal (highlighted in yellow) modulates specifics weights. Here, we show only those weights that involve the third neuron in the network, but the same patterns of weights apply to all neurons in the network. WTA: winner-take-all. **c)** Example signals in each layer of the network, shown for two successive decisions (trip 1 in black, and trip 2 in gray). The final output in the 5th layer selects the best shortest trip, given the expectations about the environment encoded in the input weights 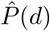 and 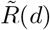. We assume that the forager executes the best shortest trip, and updates the weights *P* (*d*) accordingly. Shown for 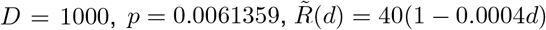, and 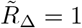.

- **Layer 1:** encodes signals about expected properties in the environment
- **Layer 2** [receives inputs from Layer 1]: encodes the expected benefit (i.e., increase in energy rate) of encountering a new food patch at each location during foraging relative to staying on the original food patch and relative to traveling
- **Layer 3** [receives inputs from Layers 1 and 2]: computes the relative benefit versus cost of taking a single trip of any length, and uses the rectify and winner-take-all operations to identify the best single trip, if it exists.
- **Layer 4** [receives inputs from Layers 1, 2 and 3]: computes the relative benefit versus cost of taking a shorter trip given the best single trip, and uses the rectify and winner-take-all operations to identify the best shorter trip (if it exists)
- **Layer 5** [receives inputs from Layers 3 and 4]: selects the desired next trip

We assume that the activity in the first layer is encoded by two sets of input weights: the first set encodes the expected density of food patches 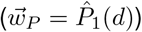, and changes over short timescales as the forager actively explores its environment. The second set encodes the expected energetic return of the environment per unit time (relative to the energy rate provided by the original food patch) 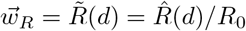, and changes over longer timescales as the animal learns about the broader environmental statistics.

The remaining computations in layers 2-5 can be performed using a combination of fixed connection weights between layers and fixed operations within layers (Fig 4b). We assume that some weights can be modulated by a global signal that encodes the cost of traveling relative to the energetic gain from harvesting the original food patch, 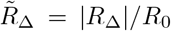 (yellow arrows in Fig 4b). All required weights and operations can be derived directly from the equations that specify the local decision rule (see SI Section 3). These weights impose an ordering over nodes in each layer, such that the activations within each layer can be interpreted as varying as a function of distance from the current food patch. As a result, given a binary input that signals when the forager is on a food patch, the network activity will return the distance that the forager should travel when searching for other food patches (Fig 4c, S1).

## Discussion

In this work, we investigated the optimal search strategy for an energetically-constrained forager that has already discovered a food patch but remains uncertain about the existence of additional patches in the environment. A key assumption of our model is that the forager remembers the location of the known food patch, and reliably returns to it in the event that no new resources are found during exploration. This assumption is motivated by the fact that many animals, including insects such as bees and ants, use internal cues to track their position relative to a reference point [10, 14–16]. Such path integration allows them to explore outward from a food source while maintaining the ability to return. Ants, in particular, are well known for their ability to return home along a direct path, even after long and circuitous explorations [16]. Crucially, this assumption of spatial memory distinguishes our framework from the classical Marginal Value Theorem, which typically assumes that a forager departs from one patch and conducts a stochastic, memoryless search for the next one [1, 8]. In that setting, the decision to leave is framed as a trade-off between diminishing returns of the current patch and the expected gain from randomly encountered new patches, with no guarantee of return even if no other patches are available.

Other studies of optimal foraging that do take into account spatial memory often posit that the forager knows the locations of all available resources. In such settings, the problem reduces to finding the most efficient route through a set of known targets, often framed as a traveling salesman problem. This formulation has been used to model traplining behavior in pollinators such as bumblebees, which develop stable visitation sequences to known flowers after repeated experience [17,18]. Similar behavior has been observed in hummingbirds, which establish repeatable foraging circuits among flower patches [19], and in primates, which revisit productive trees in a consistent order when foraging in familiar areas [20]. While models of such behavior address how animals exploit known environments efficiently, our work focuses on how to structure exploratory search before the spatial structure of the environment is known.

Our model provides a framework for analyzing how such decisions should be structured under environmental uncertainty and energetic constraints. By accounting for all possible states of the environment, including whether and where other patches might be present relative to the known food patch, we found that the optimal strategy depends critically on the forager’s prior expectations. When the environment is expected to be sparse, a single long exploratory trip is favored; when patch density is believed to be high and the quality of new patches is expected to decline steeply with distance, it is beneficial to break the exploration into multiple successive trips. This latter pattern, in which foragers make repeated excursions that gradually increase in length, has been observed in *Drosophila* [13]; our results offer a normative explanation for why such behavior may be beneficial. While it remains challenging to access animals’ prior beliefs, it is nevertheless possible to shape these beliefs through prior exposure to environments that differ in the density or quality of resources [21]. Our results provide concrete predictions for how behavior should differ across these scenarios.

To better understand the computational cost of optimal planning, we derived a local decision rule that approximates the optimal strategy. Although the component calculations involve integrals that may at first appear complex, we showed how these computations can be implemented in neural circuits using fixed connections between populations of neurons, with each population encoding key spatial quantities. The resulting circuit architecture is both conceptually straightforward and biologically plausible, relying on feedforward connectivity and local operations such as additive and multiplicative integration of inputs. As such, our approach aligns with efforts to hand-design biologically plausible circuits for implementing specified computations [22–24], and offers a level of mechanistic and computational interpretability that can be difficult to extract from task-optimized neural networks [25]. More broadly, our work also touches on the nature of optimal decision-making in small brains with constrained neural circuits. In many contexts, generating optimal behavior can seem computationally demanding, and might appear to require explicit deliberation, weighing of alternatives, or even conscious reasoning. Yet animals routinely make rapid and seemingly effortless decisions that govern their survival. Our results show how near-optimal strategies can emerge from neural networks that implement local, low-level operations. This suggests that evolution may endow organisms with neural architectures that effectively “solve” complex optimization problems, not through abstract reasoning but through hardwired structure and dynamics.

To make our problem tractable, we made a number of simplifying assumptions. We considered a one-dimensional environment, assumed a fixed belief about resource density, and treated the known food patch as non-depleting. We also assumed that the forager always returns successfully to a remembered location, and that it consumes sufficient energy between trips to support the next exploratory phase. While these assumptions simplify the analysis, the broader framework is flexible and can be extended to treat a broader range of scenarios. For example, trip distances could be modulated by the quality of the forager’s memory and the ability to successfully execute a return trip, to account for scenarios where the forager gets lost; similarly, feeding times could be modulated by prior exploratory bouts, to account for scenarios where the prior trip was longer or shorter than intended. If the environment is changing over time, the forager would additionally benefit from updating its beliefs about patch density and overall depletion levels based on prior experience. While such modifications introduce additional complexity, the same core approach—solving for the optimal sequence of exploratory decisions to maximize expected utility across a distribution of possible environments, and asking whether the optimal strategy can be achieved using simple rules— remains applicable. Exploring how these extensions affect the structure of the optimal strategy, and whether similarly simple decision rules can be used to approximate optimality, is an interesting direction for future work.

## Acknowledgments

This work was inspired by discussions with Hannah Haberkern, Dennis Goldschmidt, Carlos Ribeiro, and Vivek Jayaraman about the potential normative underpinnings of structured search behavior, based on nutrient- and density-dependent foraging experiments that were written up separately [13, 21]. These discussions and experiments were part of a collaboration between the Ribeiro, Jayaraman, Hermundstad, and Rubin (Gerry) labs that was supported by the Janelia Visiting Scientist Program. We thank Vivek Jayaraman, Hannah Haberkern, and members of the Jayaraman and Hermundstad labs for useful feedback on early stages of this work. YG and AMH were supported by the Howard Hughes Medical Institute.

## Supplementary Information

### 1 Expected utility of a planned trajectory

Any given planned trajectory consists of a sequence 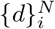 of *N* outbound trips to and return trips from locations *d*_1_, *d*_2_…*d*_*N*_ . Assuming that the forager moves at constant velocity *v* = 1 and always leaves the food patch as soon as it has enough energy for the next trip, the duration on the patch before the *i*^*th*^ trip is given by: Ω_*i*_ = 2*d*_*i*_|*R*_Δ_|*/R*_0_.

The planned trajectory is the actual trajectory that the animal takes if it does not encounter any new food spot throughout this period, which occurs with probability 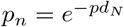. In this case, the total time the animal spends away from the original food patch is given by 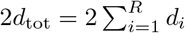, and the corresponding average reward rate *R*_mean,n_ of the animal over a time interval *T ≥* 2 (1 + |*R*_Δ_|*/R*_0_) *d*_tot_ is:

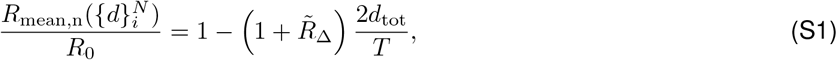

where 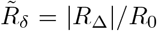. This expression captures the intuition that if no new food patch is found (which must be the case if no other food patch is present), the time spent exploring reduces the animal’s average reward rate.

If instead the forager encounters a new food patch at position *d*, we assume that the reward rate stays at *R*(*d*) from then onward (i.e., the forager stops exploring), and the mean reward rate 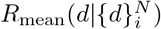 over the same time interval *T* for the planned trajectory is:

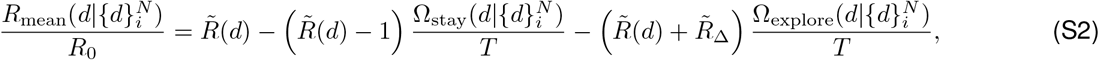

where 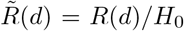, and 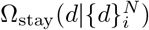 and 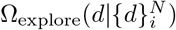 are respectively the total time the animal spends on and away from the original food patch before encountering the new food patch.

For any planned trajectory 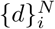, the overall average mean reward rate ⟨*R*_mean_ ⟩, with ⟨ · ⟩ representing the average over the distribution of environments (i.e., potential locations of the nearest food patch), is then given by:

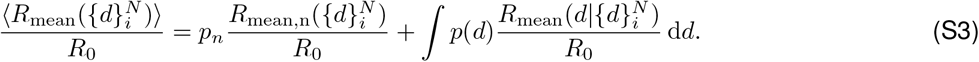

To obtain the optimal planned trajectory with *N* trips, we solve for the set of trips 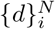 that maximizes ⟨*R*_mean_ ⟩.

### 2 Environment with sparse patches and uniform nutritional content

Here, we assume that the nutritional content of patches is constant across space 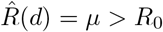, and that patches are sparse. In this limit of *pD «* 1, the probability that the nearest patch is at location *d* is given by *P*_1_(*d*) = *pe*^*−pd*^ *≈p*.

When comparing the utility of taking 2 trips instead of the best single trip, the loss in utility from the scenario where the second patch is far (scenario 2) is given by (Eq. 2):

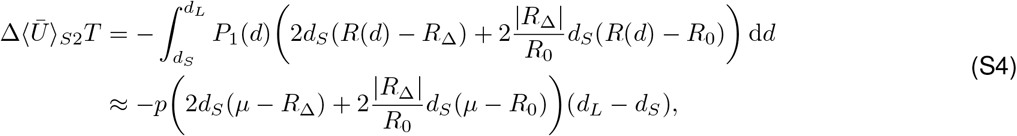

while the gain in utility from the scenario where the second patch is near (scenario 3) is given by (Eq. 3):

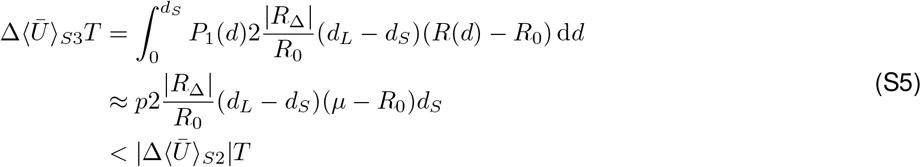

### 3 Possible neural circuit implementation of local decision rule

We consider a neural network architecture where there is an indicator neuron that is active when the forager is on the original food patch (and hence has to decide whether and when to leave the food patch and how far to travel on its next trip). The activity of this neuron (*s*^*I*^ = 1 when active and *s*^*I*^ = 0 when inactive) acts as an input to three sets of neurons in Layer 1 (Fig S1a):

- *P* **- neurons:** A set of *N* neurons with input weights 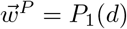 representing the probability distribution for the location of the nearest patch.
- 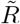**- neurons:** A set of *N* neurons with input weights 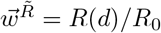 representing the expected energy rate from a potential new patch relative to that from the current patch.
- 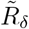 **-neuron:** A single neuron with input weights 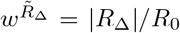, representing the ratio between the energy rate expenditure from traveling and the energy rate gain from the current patch.

The belief about the environment, including both the availability of new patches as well as the utility from them, is stored in the weights 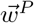 and 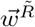, and we assume that these can change over time. In particular, 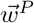 should be updated as the animal explores and learn about its environment (absence/presence of new food spot), while we expect 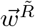 and 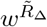 to be changing over longer timescales. Since the activity of the indicator neuron is binary, the activities for the different neuron types 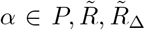, are also the corresponding input weights when the indicator neuron is active (Fig S1a):

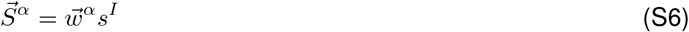

With this, we then ask how the output activities of these Layer 1 neurons can serve as inputs to downstream neurons to determine the desired next trip length and the feeding duration on the current patch before leaving.

#### 3.1 How far to travel

Recall that the two-step local rule for determining whether to leave the food patch, and how far to travel on each successive trip is as follows:

**Step 1:** Determine the optimal length of a single trip, 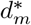. If the expected utility of this trip is greater than that of staying on the original food patch, proceed to Step 2; otherwise, stay on the original patch and continue feeding.

**Step 2:** Weigh the expected benefit and cost of first taking a shorter trip to 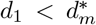 (assuming that if no new patch is found on the trip to 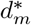, the forager continues exploring the remaining space), and determine the optimal value of 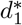 that maximizes the benefit minus the cost. If the benefit of taking 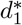 exceeds the cost, plan to travel to 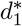; otherwise, plan to travel to 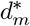.

To construct a circuit that takes inputs from the Layer 1 neurons and outputs a representation of the next trip distance (either 0 or 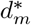 or 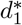 ), it is therefore useful to break down the computations that are required for each step.

**Step 1:**

From Eq. S3, the relative advantage of making a single-trip trajectory to *d*_*m*_ compared to remaining on the food original patch for a period of time *T* is given by:

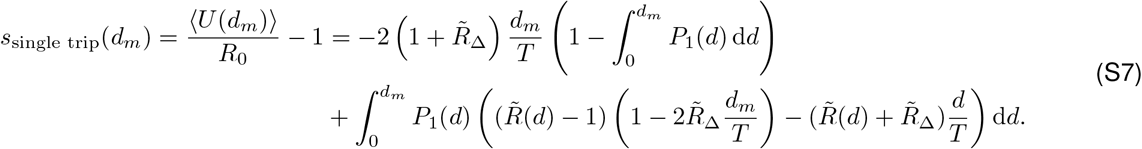

To carry out this computation, it is useful to have two sets of *N* neurons in a downstream layer (Layer 2), the activities of which are given by (Fig S1b):

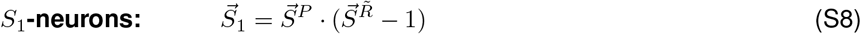

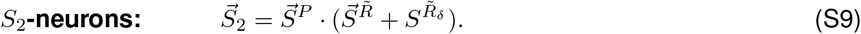

This can be achieved if each neuron *i* = 1, 2, …, *N* in these sets receives inputs from the corresponding *i*^*th*^ *P* - and 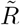- neurons, as well as from the 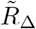 neuron in the case of the *S*_2_- neurons. The signal from the 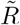 - neuron is either offset by 1 (for the *S*_1_- neurons) or summed with the activity of the 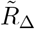 neuron (for the *S*_2_- neurons), before combining multiplicatively with the activity from the *P* -neuron.

We then consider another set of *N* neurons (‘*d*_*m*_- neurons’) in a downstream layer (Layer 3) that receives inputs from both Layer 1 (*P* - and 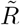- neurons) and Layer 2 (*S*_1_-, *S*_2_- neurons), such that the overall external inputs 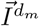 to these *d*_*m*_- neurons are given by (Fig S1b):

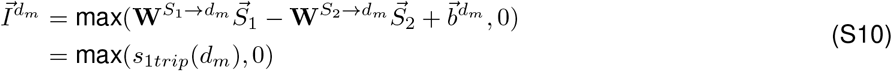

where the weights 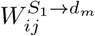 are the connection strengths from the *j*^*th*^ *S*_1_-neuron to *i*^*th*^ *d*_*m*_-neuron, 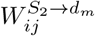 are the connection strengths from the *j*^*th*^ *S*_2_-neuron to *i*^*th*^ *d*_*m*_-neuron, and 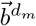 is the bias term that depends only on inputs from Layer 1 (namely, the *P* - and 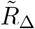- neurons).

Based on the expression for *s*_1*trip*_(*d*_*m*_) (Eqn. S7), the elements of these weights and bias terms are given by:

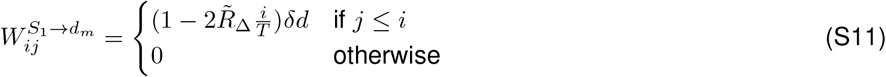

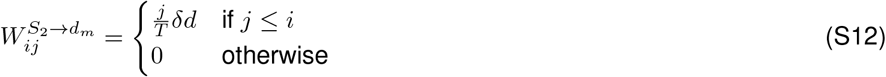

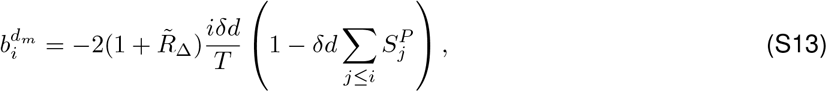

where *δd* is the spatial resolution. It is useful to note that the structure of these input connections is fixed and encodes the spatial geometry and structure of the problem. For some of these connections (namely those from the *S*_1_- and *P* - neurons), their connection strengths are modulated by 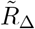, suggesting that the activity of the 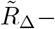 neuron serves as a global signal for the whole network. Therefore, it might also be useful for the value of 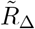 to be encoded by certain chemical/hormone levels, and one can imagine the activity of the 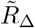-neuron regulating the release of such chemicals/hormones.

One could then imagine additional connections among the *d*_*m*_- neurons that implement a winner-take-all operation, such that only the neuron that receives the highest input will remain active (Fig S1b), and the identity of this active neuron encodes 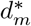. In other words, the activities 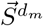 of these neurons are given by:

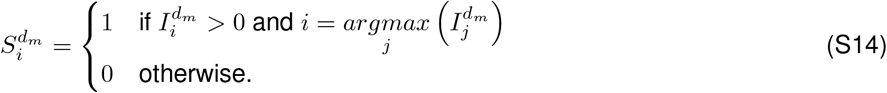

This representation of 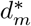 can then be used to determine whether it is advantageous to take a shorter trip first.

**Step 2:**

Analogous to the *d*_*m*_- neurons that provide a representation of the value of 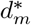, we assume that there exists another set of *N* neurons(‘*d*_1_*−* neurons’) whose binary activities 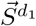 encodes the best shorter trip 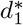 (if it exists). This can be achieved if the net inputs 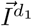 to these neurons are given by the net benefit of traveling to *d*_1_ (if positive, and zero otherwise), and additional connections among the *d*_1_- neurons implement a winner-take-all operation that result in at most one of those neurons being active (Fig S1c):

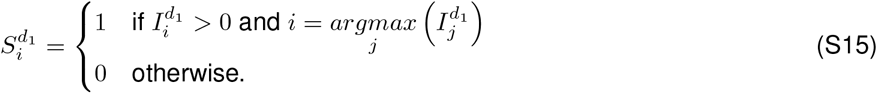

To infer the network connections (between the upstream neurons in Layers 1-3 and these *d*_1_- neurons) needed to achieve the desired inputs 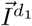, we return to the computations required.

From Eqs. 1-3, and assuming that the forager eventually explores the whole space (of length *D*) if no new patches are found, the net benefit (expected gain in average energy rate) of first taking a short trip to 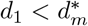 relative to the baseline energy rate of staying on the current food patch is given by:

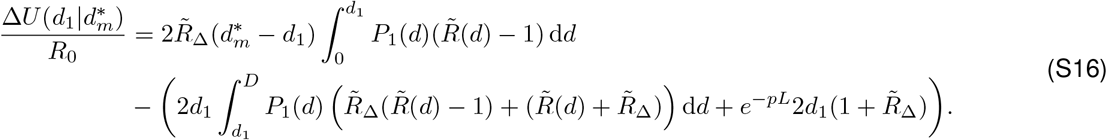

The desired inputs to the *d*_1_- neurons can therefore be expressed as:

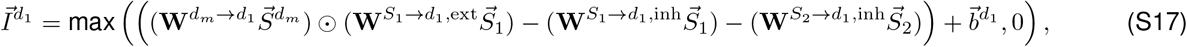

where the weights from the *d*_*m*_ neurons 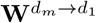 are:

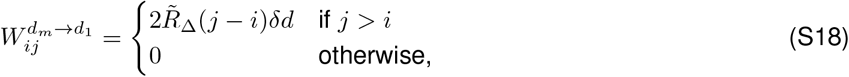

the weights of the activating connections 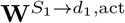 from the *S*_1_*−* neurons are:

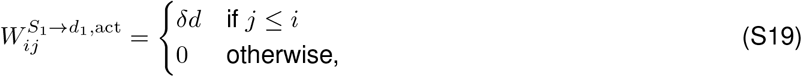

the weights of the inhibitory connections 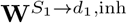 from the *S*_1_*−* neurons are:

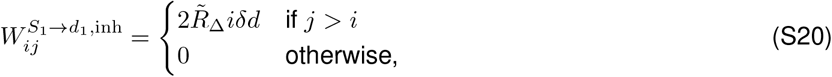

the weights of the inhibitory connections 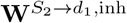 from the *S*_2_*−* neurons are:

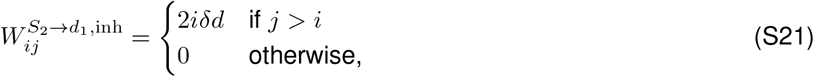

and the bias terms 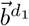 (that depend only on inputs from Layer 1) are given by:

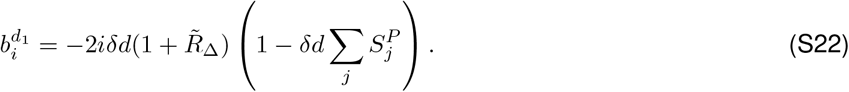

As for the input connections to the *d*_*m*_- neurons, these input connections to the *d*_1_- neurons also encode the spatial geometry and structure of the problem, with some of the weights modulated either directly or indirectly by the 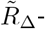 neuron.

Finally, a set of *N* neurons (‘*d*_next_- neurons’) receives inputs from both the *d*_*m*_ and *d*_1_ neurons to provide a representation of the desired next trip distance (Fig S1d). The input weights to the *i*^*th*^ *d*_next_-neuron are given by:

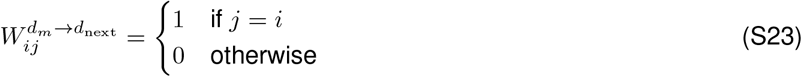

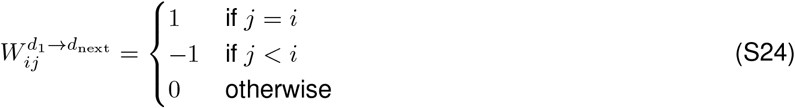

such that the activities of the *d*_next_- neurons are given by:

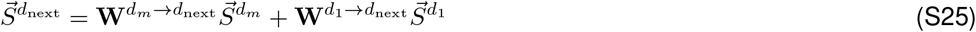

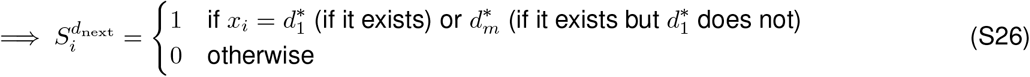

#### 3.2 How long to stay on food patch before leaving

The set of *d*_next_- neurons that encode how far the forager wants to go on its next trip and another set of neurons (e.g. place cells) that encode the current location of the forager, together with the global signal 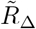, can serve as inputs to a downstream neuron to compute the duration on the food patch before leaving 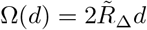.

**Figure S1:**
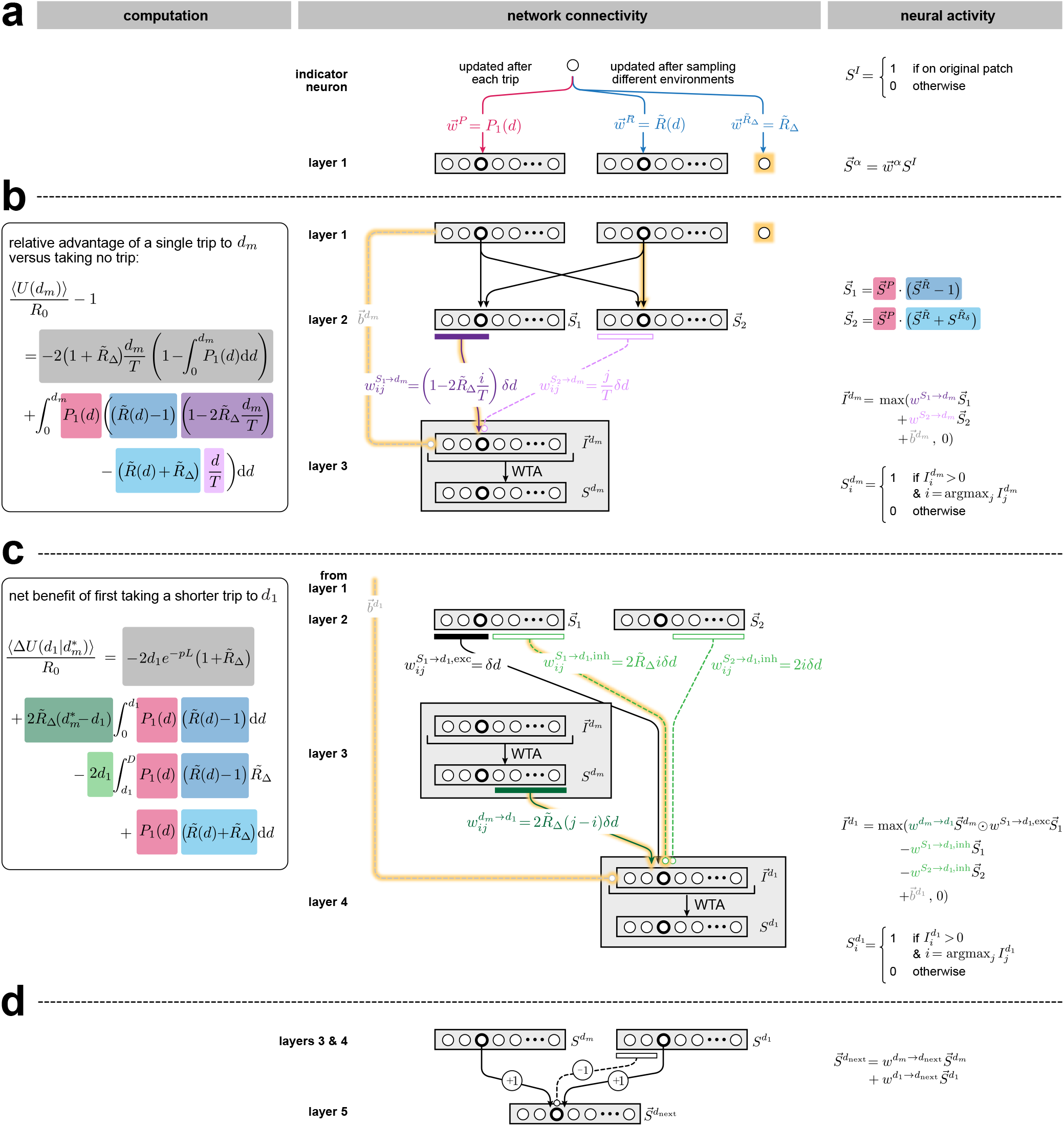
Breakdown of neural network that implements the local decision rule. **a)** Schematic of an indicator neuron (representing whether the forager is on the original food patch) and how it is connected to Layer 1 neurons that encode properties of the environment. **b)** Schematic showing how computations of the relative advantage of a single trip compared to remaining on the original food patch (step 1 of the local rule) can be implemented in the circuit using additional layers of neurons. **c)** Schematic showing how computations of the net benefit of taking a shorter trip first (step 2 of the local rule) can be implemented in the circuit using an additional set of *d*_1_ neurons that encodes the best shorter trip. **d)** Schematic showing how *d*_*m*_ and *d*_1_ neurons can provide inputs to a final set of *d*_next_ neurons to provide a representation of how far the forager should go next.

